# Transcriptome Variant Analysis of Noise Susceptibility in C57BL/6J Mice

**DOI:** 10.1101/2022.07.11.499624

**Authors:** Siyue Wang, Jing Cai, Ligang Kong, Lei Xu, Zhaoming Fan, Haibo Wang

## Abstract

**Background:** Susceptibility to noise varies dramatically between mice of the same genetic background; however, the underlying molecular mechanism remains unknown.

**Methods:** C57BL/6J (B6) mice of the same sex, age, and strain were exposed to noise of the same intensity and duration, and the auditory brainstem response (ABR) threshold was determined 48 h later. Some mice had significant hearing loss, while some did not; the ABR threshold measured in these two groups of mice was significantly different. The cochlea of the two groups of mice was dissected, and RNA sequencing and analysis were performed. Differentially expressed genes (DEGs) between the two groups were selected, Kyoto Encyclopedia of Genes and Genomes pathway analysis was performed, and protein–protein interaction network maps were listed.

**Results:** This study showed that noise exposure of the same intensity and duration caused different degrees of hearing loss in C57BL/6J (B6) mice. This was the result of the up-regulation or down-regulation of many genes, such as Nop2, Bysl, Rrp9, Spsb1, Fbxl20, and Fbxo31. Changes in the transcriptome of these genes may affect cochlear susceptibility to noise.

**Conclusion:** The DEGs identified in this experiment may provide more insight into protocols for gene therapy in the clinical practice of hearing loss.

## 1. Introduction

Noise-induced hearing loss is a form of sensorineural deafness that results from prolonged exposure to a noisy environment and a combination of factors. At present, the recognized mechanisms are mechanical damage to the cochlea, metabolic damage, immune and inflammatory damage, and genetics (Ding et al. 2019). Current studies have found that the genes involved in noise-induced hearing loss are associated with oxidative stress, DNA repair, gap junctions, apoptosis, K^+^ recycling, and heat shock proteins (Ding et al. 2019; Mao and plasticity 2021; Sliwinska-Kowalska and research 2013). It is well known that susceptibility to noise varies significantly among individuals, and not everyone experiences the same hearing loss after the same noise exposure (Sliwinska-Kowalska and research 2013).

Prior to this, some animal studies have demonstrated that NOX3, FOXO3, NRF2, CX26, CRFR2, BHMT, A1AR, and MYH14 (Beaulac et al. 2021; Fu et al. 2016; Graham et al. 2010; Honkura et al. 2016; Lavinsky et al. 2015; Partearroyo et al. 2019; Vlajkovic et al. 2017; Zhou et al. 2016) knockout mice are more sensitive to noise than wild-type mice. These studies in knockout mice have shown that genetic defects in mice, which disturb specific paths and structures within the cochlea, make mice more sensitive to noise (Le et al. 2017). There are also many other studies of genetic mouse noise susceptibility (Fairfield et al. 2005; Holme and JARO 2004; Kozel et al. 2002; Ohlemiller et al. 1999; Ohlemiller et al. 2000; Schick et al. 2004; Tabuchi et al. 2005; Yan et al. 2013), and it has been demonstrated that noise susceptibility varies for different strains of mice. For example, B6 and 129 mice showed differences in gene expression after noise exposure; HSP70, HSP40, GADD45b, and P21Cip1 were significantly induced and up-regulated at the protein level in 129 mice, and their up-regulation may have a protective effect on hearing in these mice (Gratton et al. 2011). Inbred C57BL/6J (B6) mice are more likely to acquire noise-induced hearing loss than inbred CBA/Cal (CB) mice (oto-laryngologica 1992; Shone et al. 1991) because the AHL gene is reported to influence susceptibility to noise-induced hearing loss (Davis et al. 2001; Erway et al. 1996; Harding et al. 2005).

Many studies have shown that people working in environments with similar noise levels often show varying degrees of hearing loss (Henderson et al. 1993), especially in occupational noise exposure. The noise susceptibility of this population is more pronounced, with approximately 33% showing noise-induced hearing impairment and 16% showing substantial hearing impairment (Themann and America 2019). In China, the prevalence of occupational noise-induced hearing loss is 21.3% (Zhou et al. 2020); therefore, not all people exposed to noise suffer from noise-induced hearing loss. In addition, many studies have employed single nucleotide polymorphism screening methods in noise-exposed populations (Liu et al. 2021; Miao et al. 2019; Zhang et al. 2019b; Zhang et al. 2019c), demonstrating that genes play an important role in noise susceptibility.

We have also found that the same batch of mice showed different degrees of hearing loss under the same noise exposure. Some mice show severe deafness immediately after noise exposure, while some do not. However, there is no study on the transcriptome of mice of the same strain with different susceptibilities to noise. Therefore, we investigated the transcriptome variation of inbred C57BL/6J (B6) mice with different noise susceptibility.

RNASeq is the first sequencing-based method to detect the entire transcriptome in a high-throughput and quantitative manner, and it can accurately quantify the expression levels of genes (Marioni et al. 2008; Wang et al. 2009b). In recent years, the advent of RNASeq technology has allowed us to discover new genes and transcriptomes for a wide range of diseases, contributing to the discovery of disease-causing factors.

Therefore, we established a noise-induced hearing loss mouse model. After noise exposure, the degree of hearing loss was determined by auditory brainstem response (ABR) measurement, and some mice were selected as the noise-resistant group (R_NE) and some as the noise-sensitive group (S_NE). The mouse cochleae were collected, and using RNASeq technology, differentially expressed genes (DEGs) related to noise susceptibility were selected. The functions of these differential genes were summarized by Kyoto Encyclopedia of Genes and Genomes (KEGG) pathway analysis in order to determine their roles in the etiopathology of noise-induced hearing loss for future studies.

## 2. Materials and Methods

### 2.1. Mice

Fifty inbred C57BL/6J (B6) male 8-week-old normal hearing mice (all purchased from Pengyue Company, Jinan, China) were selected and maintained in a quiet environment in the specific pathogen-free animal room of the Shandong Institute of Otolaryngology. All mice lived in a room with constant temperature (approximately 22–25°C) and were given adequate food and water. One week later, 40 mice were randomly selected as the experimental group and exposed to noise to establish the noise-induced hearing loss model, and 10 mice were used as the control group without noise exposure (Control). According to the hearing results after noise exposure, we further subdivided the experimental group: those with a hearing threshold range above one standard deviation of the mean hearing threshold of the experimental group comprised the S_NE, and those with a hearing threshold range below one standard deviation of the mean hearing threshold of the experimental group comprised the R_NE.

All animal experiments were approved by the Ethics Committee of the Shandong Provincial ENT Hospital, Shandong University, and the experiments complied with the relevant ethical regulations for animal testing and research. All efforts were made to minimize the number of animals used and to prevent their suffering.

### 2.2. Noise exposure

Mice were anesthetized intraperitoneally with a mixture of ketamine hydrochloride (100 mg/kg) and xylazine (4 mg/kg) and then exposed to 100 dB sound pressure level (SPL) white noise for 2 h. The noise stimuli were synthesized by a noise generator (SF-06, Random Noise Generator, RION, USA) and amplified by an amplifier (CDi 1000 Power Amplifier, Crown, USA). The bottom of the cage and the center of the speaker were placed on the same horizontal line, and the distance between the two was determined using a noise meter. The noise meter radio was placed in the center of the bottom of the cage, and the noise measured in the center was ensured to be 100 dB each time.

### 2.3. ABR measurement

All mice were tested for ABR thresholds prior to the experiment, and the experimental group underwent re-testing for ABR 48 h after noise exposure. The anesthesia employed was the same as above, and the anesthetized mice were placed in a sound-proof chamber to measure ABR responses under sound stimuli at 4, 8, 12, 16, 24, and 32 kHz, with 1024 repetitions of stimulation per recording (Tucker-Davis Technology, USA). The left ear of the mouse was oriented towards the speaker (MF1; TDT) at a distance of approximately 5 cm, and the recording electrode was inserted into the subcutaneous tissue of the middle of the two ears, the reference electrode was fixed at the ipsilateral ear, and the ground electrode was placed at the back. The sound level was decreased by 5 dB from 90 dB until no hearing curve appeared. We ensured that each frequency was judged by the same person and that reliable results were obtained. It is also essential to duplicate this operation for low SPLs close to the threshold to guarantee the stability of the signal. After ABR audiometry, the mice were laid on a warming mat to maintain body temperature and ensure awakening.

### 2.4. Sample collection and preparation

ABR data were recorded for comparison, and the cochleae of mice in the R_NE, the S_NE, and the Control were collected. First, the mice were anesthetized with the same drug dose and methods, and cardiac perfusion was performed using normal saline. The left and right cochleae were removed and rapidly placed in RNA later (Invitrogen, AM7021) overnight at 4°C and transferred to -20°C for long-term storage. Total RNA was extracted and tested for quality. Polymerase chain reaction amplification was performed to complete the entire library preparation work.

### 2.5. RNA sequencing

Qualified libraries were sequenced with an Illumina NovaSeq6000 sequencer with a sequencing strategy of PE150 to obtain high-quality sequences (Clean Reads).

### 2.6. DEG analysis

DESeq2 (1.20.0, method = ‘per-condition’) was used for gene differential expression analysis. Differential gene screening primarily means the fold difference (fold change value) and q value (padj value, corrected P value) are related metrics. The criteria for differential gene screening in this experiment were more than 1.5-fold difference and q < 0.05.

### 2.7. Protein–protein interaction network maps

Using the STRING protein interaction database, combined with the results of DEG analysis and the interaction pairs included in the database, the DEG sets can be directly mapped to the protein–protein interaction (PPI) network of this species.

### 2.8. KEGG pathway analysis

KEGG is a record base for the systematized analysis of genome functions that link genomic and higher-order functional information. Pathway analysis was performed by applying a hypergeometric test to each pathway in KEGG to identify pathways that were evidently enriched in DEGs.

## 3. Results

### 3.1. Detection of ABR thresholds in mice before and after noise exposure

To compare the differences in hearing of mice before and after noise exposure, we measured the ABR threshold of all mice before noise exposure, followed by noise exposure immediately after in the experimental group, and then measured the ABR threshold of noise in mice 48 h later; the data were then compared (Figure 1). According to the hearing results after noise exposure, we screened out the R_NE and the S_NE. The susceptibility of C57BL/6J (B6) mice to noise showed individual differences under the same noise exposure.

**Figure 1.**
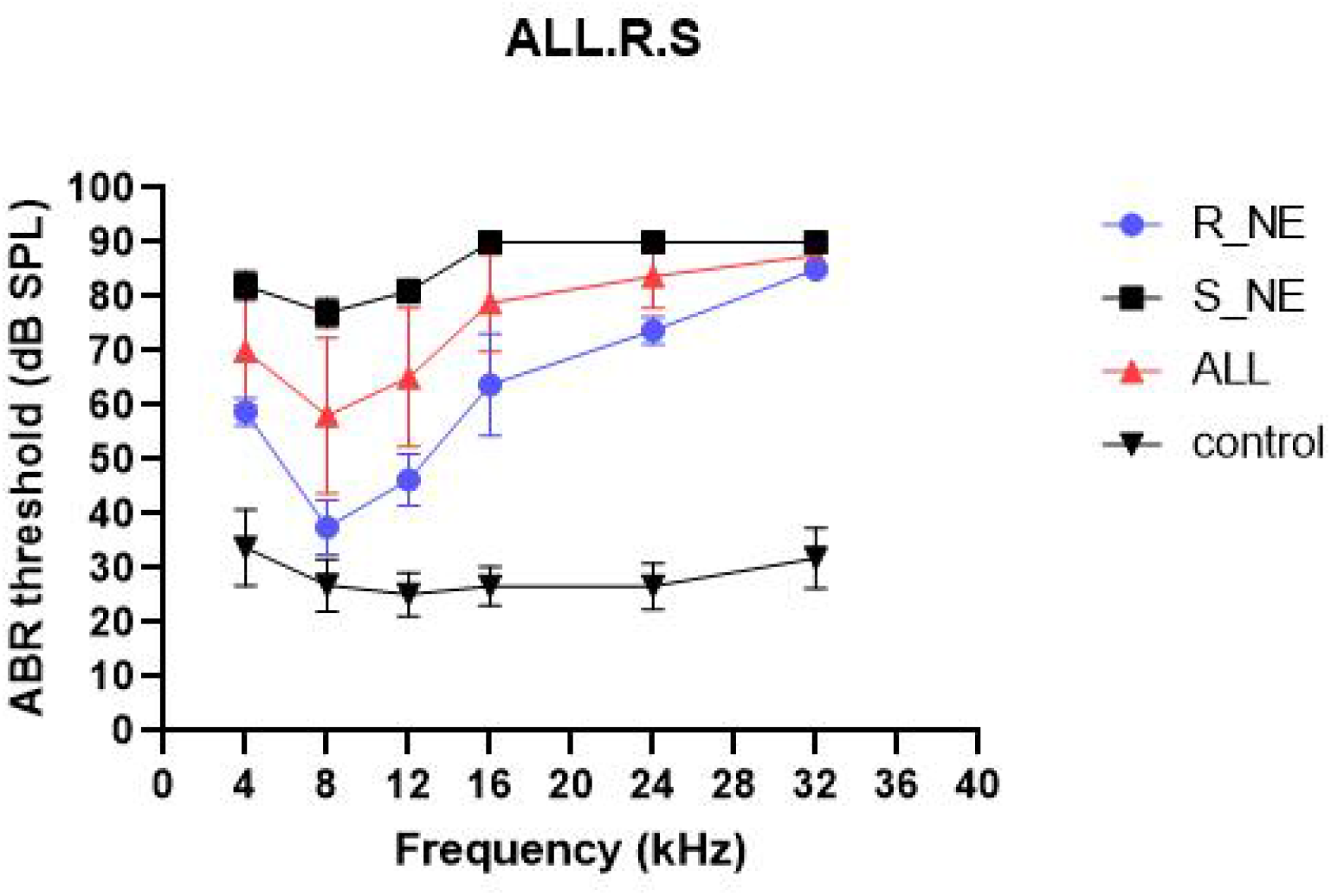
Determination of auditory brainstem response threshold in the three groups of mice The R_NE is the noise-resistant group, the S_NE is the noise-sensitive group, the NE is the experimental group, and the control is the control group without noise.

### 3.2. Gene expression in mouse cochleae with different susceptibility to noise

C57BL/6J (B6) mice showed different hearing loss under the same noise exposure. To compare the differences in gene expression among the three groups (the R_NE, the S_NE, and the Control), we performed principal component analysis of RNASeq data, estimated the PC1 and PC2 values of each sample, and plotted the results (Figure 2). The gene expression analysis of the three groups cluster together and converge into three parts, indicating that there is a significant difference in gene expression between the R_NE, the S_NE, and the Control.

**Figure 2.**
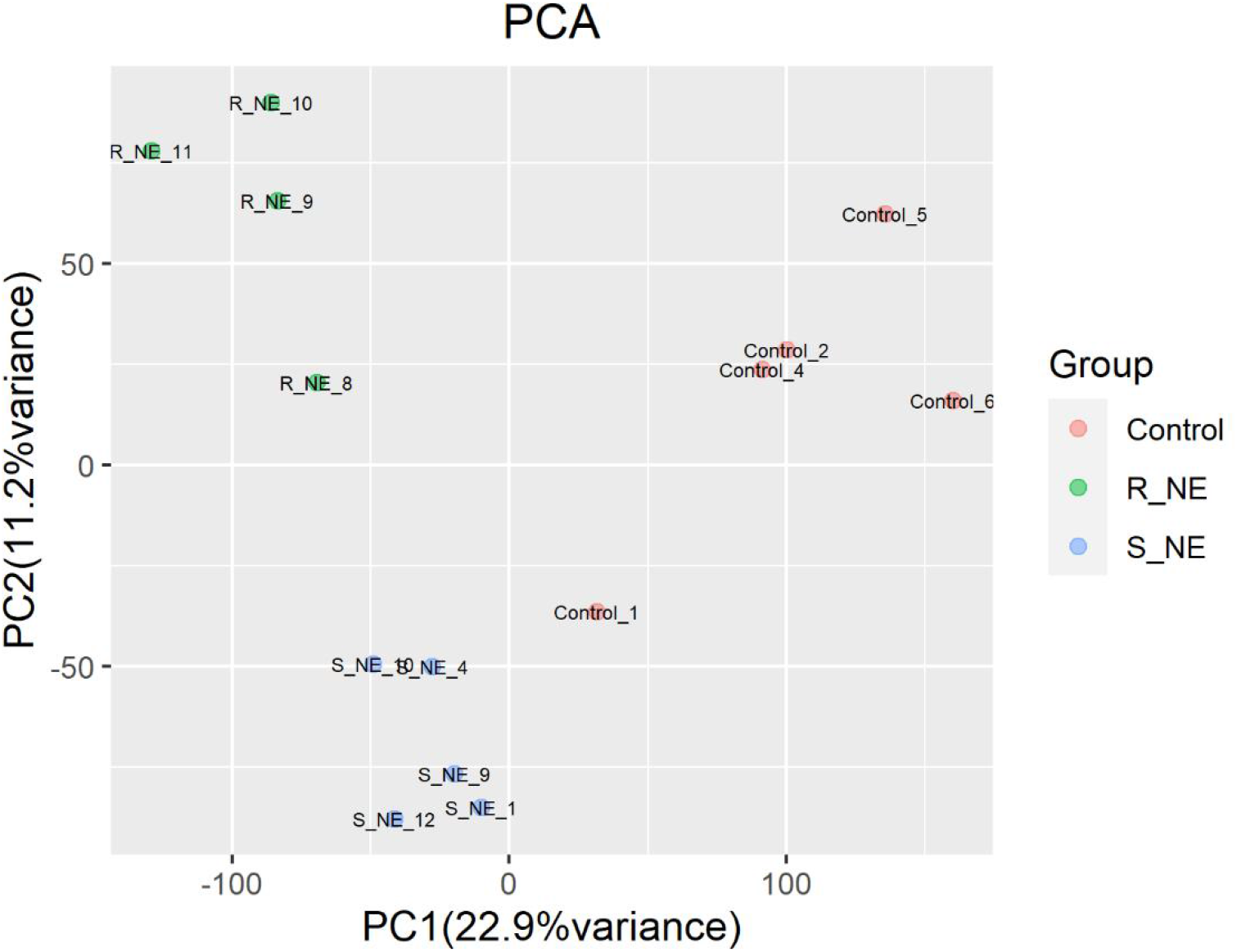
Principal component analysis Red dots are the Control, green dots are the R_NE, and blue dots are the S_NE. The three groups of samples are clustered into three parts, which indicates that there are significant differences in gene expression in the cochlear samples.

Differential analysis was performed between the three groups; the differential genes were plotted on a Wayne diagram (Figure 3). In total, there were 802 differential genes, 559 up-regulated and 243 down-regulated, in the S_NE compared to the Control. Further, there were 2646 differential genes, 1576 up-regulated and 1070 down-regulated, in the R_NE compared to the Control. These three groups shared 529 common differential genes. These 529 differential genes were all noise-induced variants and were significant within the R_SE and the S_NE.

**Figure 3.**
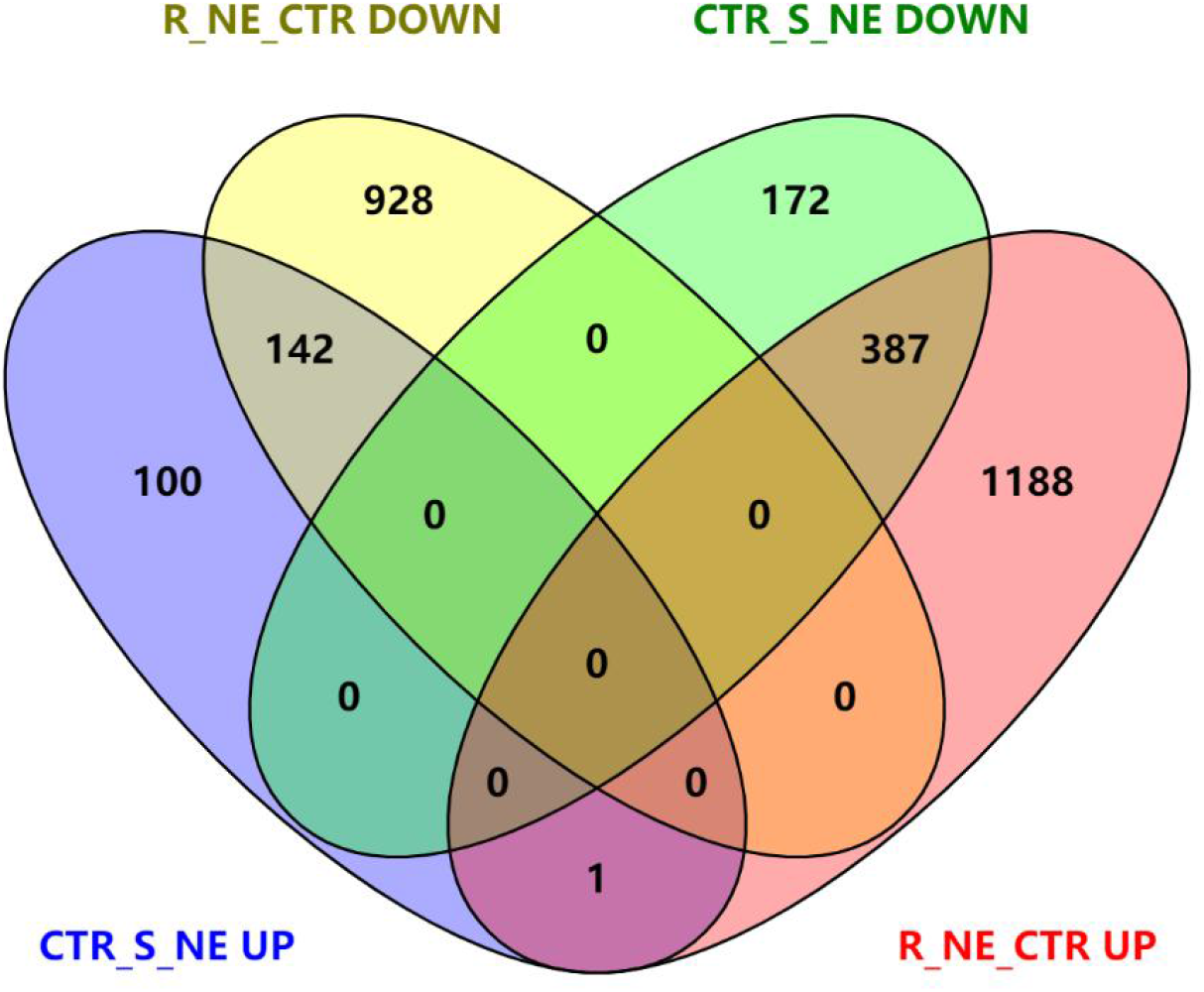
Wayne diagram

Next, we focused on the differential genes between the R_NE and the S_NE. A total of 695 differential genes were obtained from these two groups by sequence analysis, with 366 genes up-regulated and 329 genes down-regulated in the R_NE compared to the S_NE (Table S1). The 695 genes were plotted in a heat map showing the significant differences in Figure 4.

**Figure 4.**
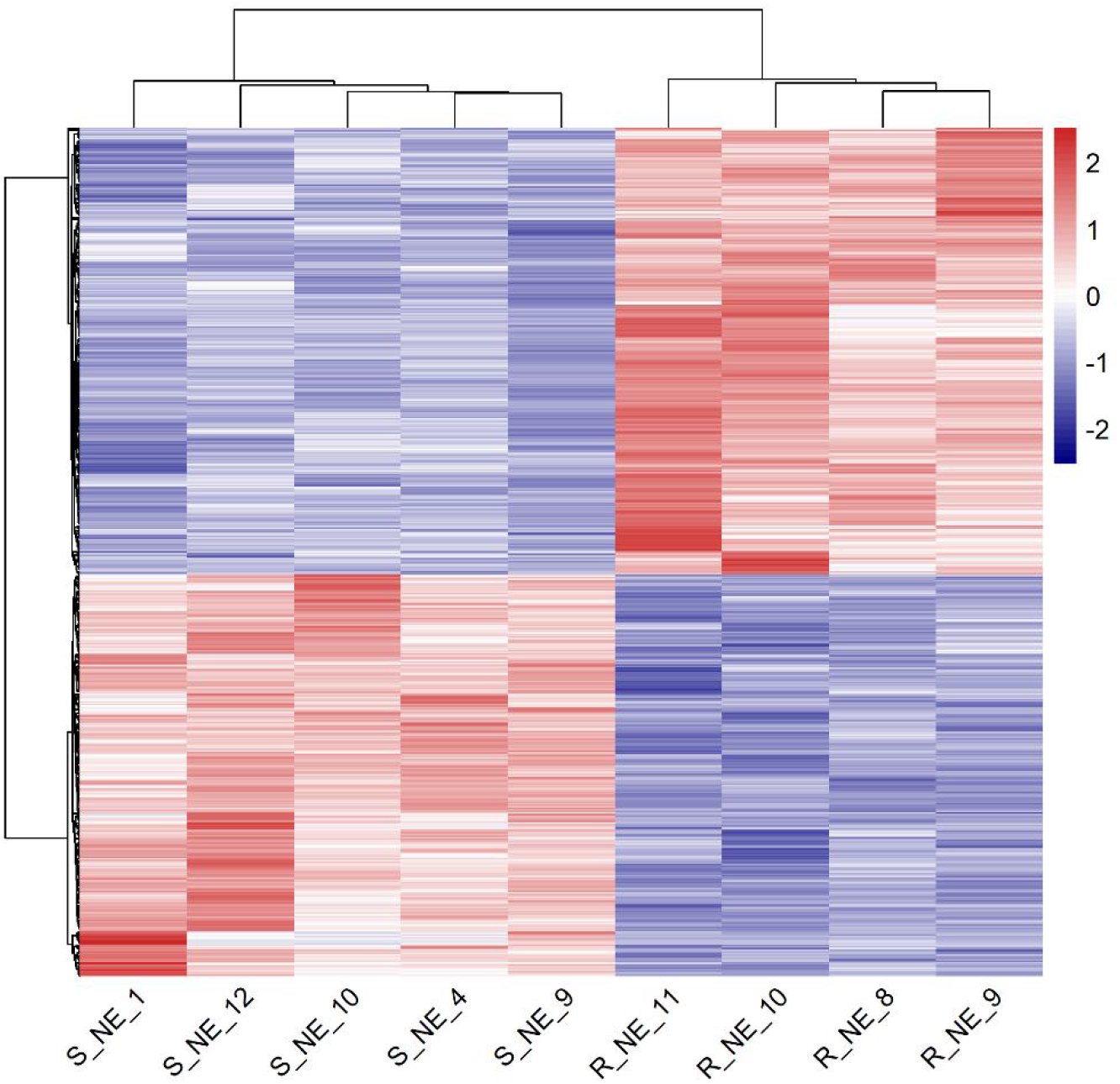
Heat map of differential genes in the R_NE and the S_NE. The red color represents high correlation, and the blue color represents low correlation. It can be seen in the figure that there are significant differences in differential genes between the R_NE and the S_NE.

### 3.3. KEGG pathway analysis of DEGs

The KEGG analysis of the 529 common differential genes in the three groups showed that the top 20 enriched pathways (Figure 5) suggested that these genes play an important role in focal adhesion, cytoskeleton, hormone synthesis, HIF-1, cellular matrix, viral infection, and Rap1. In addition, these pathways are significant in noise resistance and noise sensitivity, and it can be concluded that noise exposure leads to mutations in these 529 genes at the transcriptional level and is associated with the above pathways.

**Figure 5.**
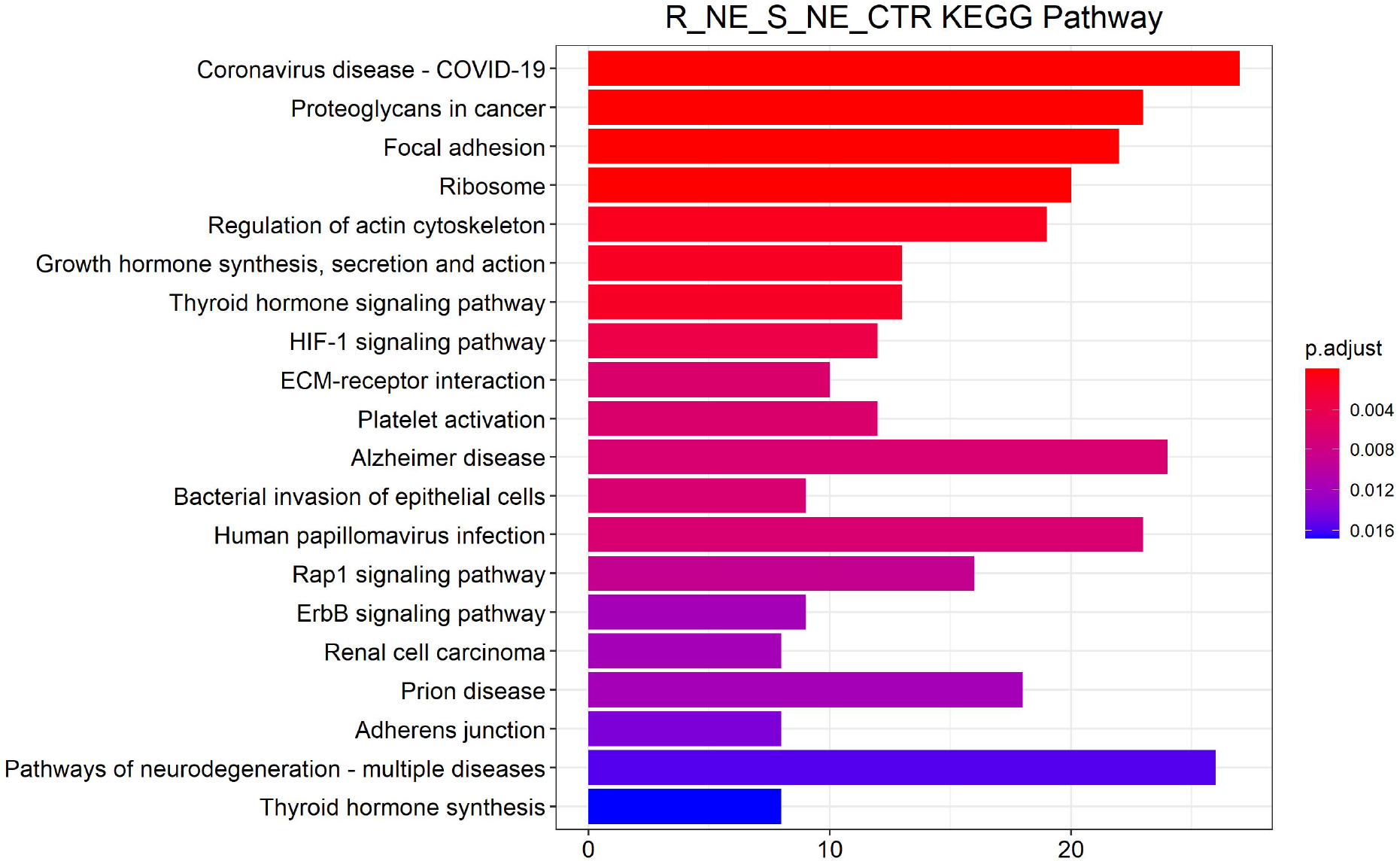
Enrichment analysis of common differential genes in the Control, the R_NE, and the S_NE

The KEGG analysis of 695 differential genes in the R_NE and the S_NE showed that these genes were significantly enriched in ribosome synthesis, apoptosis, NF-κB signaling pathway, nucleic acid metabolism, and insulin resistance pathways in eukaryotes. We believe that these pathways are significant in noise susceptibility and play a crucial part in the individual differences in hearing loss in mice after the same noise exposure.

### 3.4. PPI network of DEG protein products in the R_NE and S_NE

The 695 DEGs between the R_NE and the S_NEwere constructed into a PPI network (Figure 7), indicating that the interactions of the proteins encoded by these genes are also closely complex, and the regulation of these genes may be controlled by interactions with other members. Analysis was performed using the MCODE plugin of Cytoscape, and the top two most significant subnetworks were selected from the resulting subnetworks (Figure 8). Blue represents that the expression level of this gene was down-regulated in the R_NE, and red represents that the expression level of this gene was up-regulated in the S_NE. The obtained genes in the subnetwork are ranked according to the magnitude of the P value; the smaller the P value, the more meaningful the gene in the R_NE. Therefore, we focused on the top three down-regulated genes, Nop2, Bysl, and Rrp9, and the top three up-regulated genes, Spsb1, Fbxl20, and Fbxo31.

**Figure 6.**
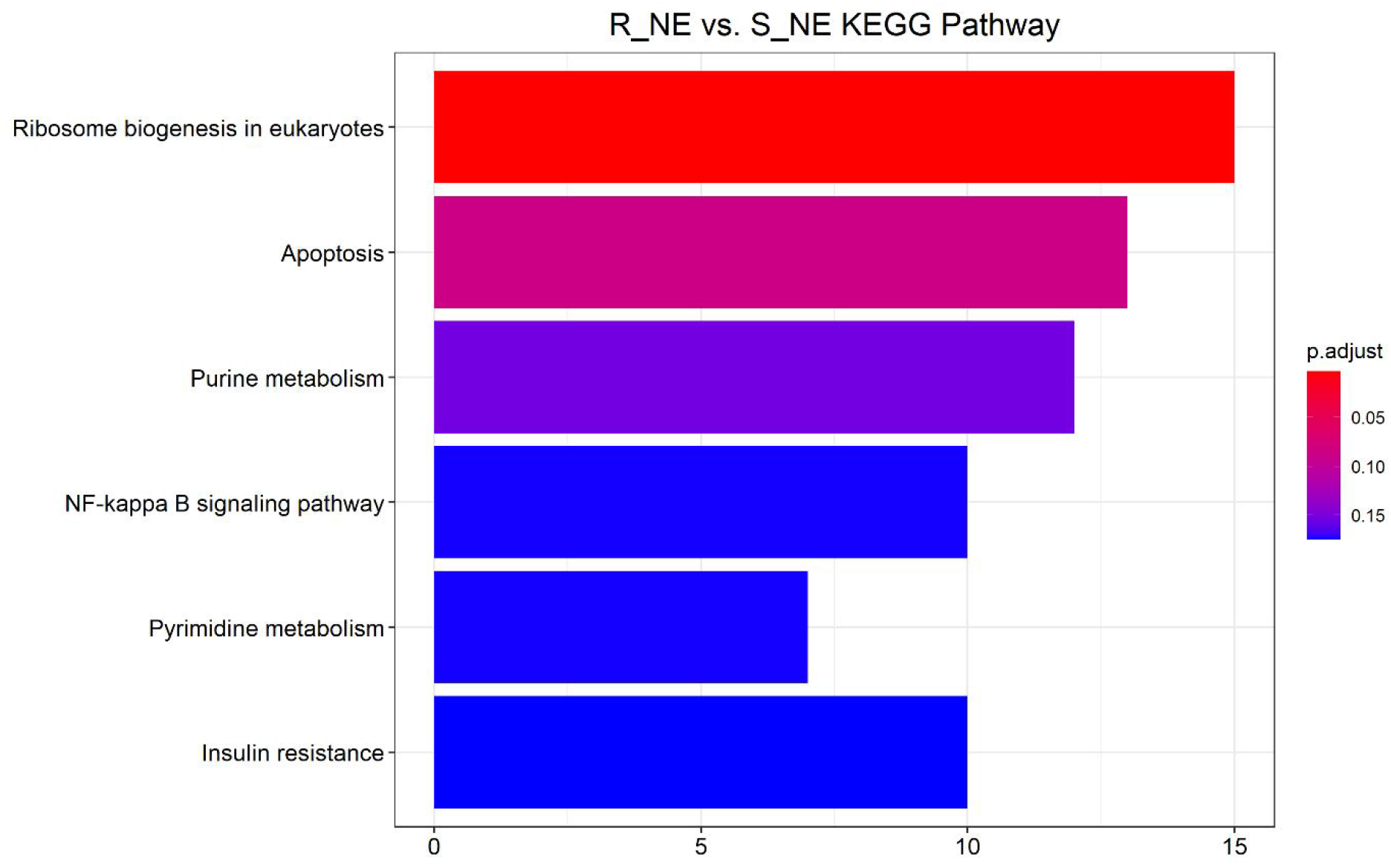
Enrichment analysis of differential genes in the R_NE and the S_NE

**Figure 7.**
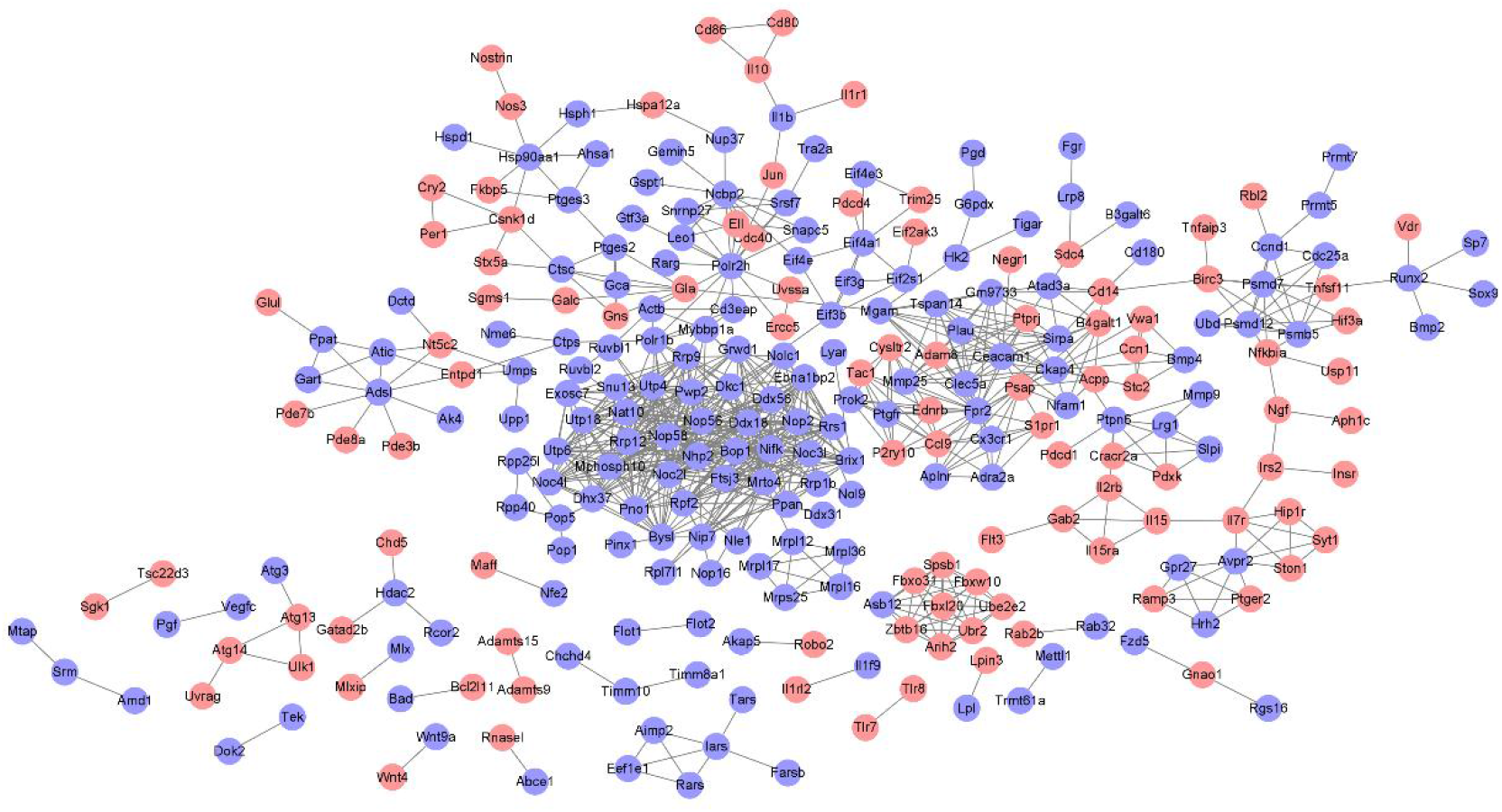
Protein-protein interaction network maps of differential genes in the R_NE and the S_NE Blue represents genes down-regulated in the R_NE, and red represents genes up-regulated in the S_NE.

**Figure 8.**
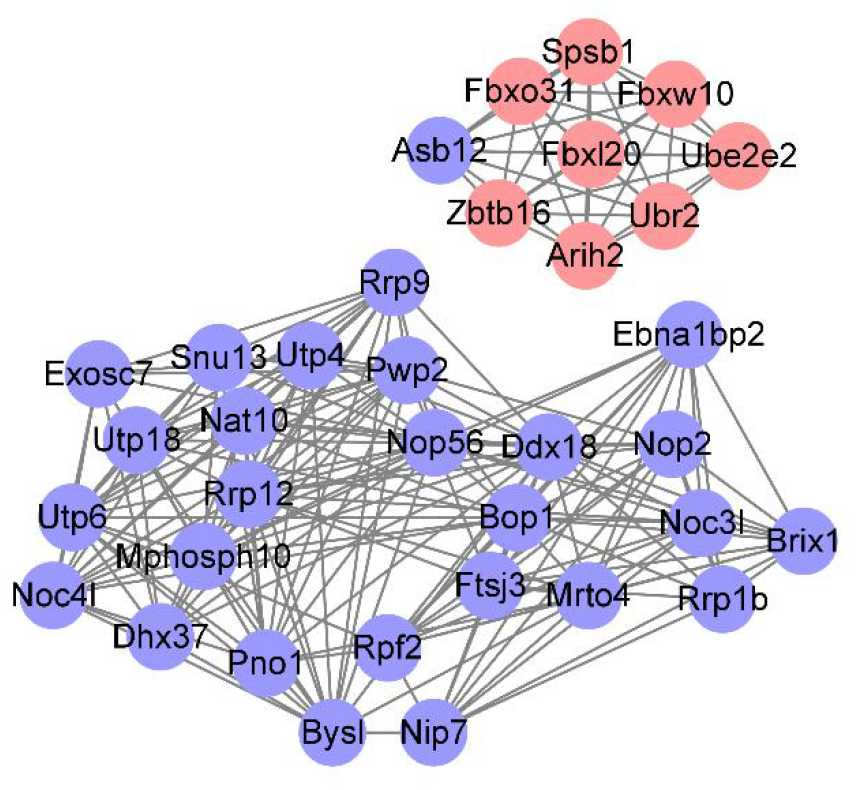
Protein-protein interaction network maps The analysis was performed using the MCODE plugin of Cytoscape, and the top two subnetworks were selected from the resulting subnetworks (upper graph). Blue represents genes down-regulated in noise resistance, and red represents genes up-regulated in noise resistance.

## 4. Discussion

### 4.1. Related DEGs for noise susceptibility

In this experiment, mice with the same genetic background showed different hearing loss due to noise exposure. Their genes were analyzed for variation arising at the transcriptional level. We found that 695 genes were differentially expressed in the R_NE and the S_NE and constructed a PPI network map (Figure 7) to screen the top two subnetworks (Figure 8) that were related to noise susceptibility in mice. Nop2, Bysl, and Rrp9 were the down-regulated genes in the R_NE, with P values in the top three in the subnetwork. Nop2 had the smallest P value among the selected subnetworks, illustrating that down-regulation of this gene was significantly associated with noise susceptibility. It has been demonstrated that the degree of methylation of NOP2 is associated with the dedifferentiation potential of postmitotic supporting cells into otic stem cells. NOP2 may therefore play a role in regulating the stemness of the organ of Corti (Waldhaus et al. 2012). We have reason to believe that NOP2 can affect cochlear susceptibility to noise; however, this needs to be confirmed in further studies. In addition, Bysl and Rrp9 are not currently being studied in the field of otology. However, some scholars have found that in liver cancer, loss of Bysl induces apoptosis (Wang et al. 2009a). Therefore, we have inferred from our experimental results that down-regulation of Bysl increases noise-induced apoptosis, resulting in cochlear sensitivity to noise; however, this requires further verification. Rrp9 is less well studied; it is a U3 snoRNA-binding protein consisting of a WD-repeat domain and an n-terminus region (Zhang et al. 2013). Rrp9 is important in the processing of pre-rRNA (Du et al. 2021), but we are currently unable to determine the effect of this gene on noise susceptibility.

Spsb1, Fbxl20, and Fbxo31 were up-regulated genes in the R_NE, with P values in the top three in the subnetwork; hence, they are also significantly correlated with noise susceptibility. Several studies have shown that up-regulation of Spsb1, Fbxl20, and Fbxo31 inhibits apoptosis in cancer (Feng et al. 2014; Kim et al. 2019; Liu et al. 2018; Manne et al. 2021; Qin and discovery 2014); hence, it is speculated that high expression of these three genes may inhibit noise-induced apoptosis and thus resist noise. We aim to continue to investigate whether up- or down-regulation of these genes protects or impairs hearing in noisy environments in subsequent experiments.

### 4.2. Pathways involved in susceptibility to noise

We further investigated the function of these differential genes. The common differential genes of the Control, the R_NE, and the S_NE were caused by noise exposure, and their pathways were associated with noise-induced hearing loss. In the present study, the pathways closely related to noise-induced hearing loss and of interest to us were the focal adhesion, regulation of actin cytoskeleton, and HIF-1 signaling pathways. The involvement of DEGs associated with noise-induced hearing loss in focal adhesion was mentioned in an article on proteomics (Miao et al. 2021). In another study, it was shown that noise can increase the expression of focal adhesion kinase, and in noise-exposed Corti organs, FAK p-Tyr577 can be detected in the outer hair cell stereocilia in noise-damaged areas (Jamesdaniel et al. 2011). A similar article also mentions focal adhesion, which highlights that nonerythroid spectrin alpha II plays a key role in the morphology and auditory function of hair cell stereocilia by modulating focal adhesion signaling (Yao et al. 2022). Stereocilia are actin-based protrusions on auditory and vestibular sensory cells that are necessary for hearing and balance. They are regulated by myosin motors, actin cross-linkers, and capping proteins (McGrath et al. 2017). Studies on the relationship between regulation of the actin cytoskeleton and noise-induced hearing loss are also ongoing. Previous reports have clearly described that F-actin cleavage occurs in the hair cells of guinea pigs and cochleae of dragon cats after noise exposure (Hu et al. 2002; Raphael and neurology 1992). Thus, it can be inferred that the signaling pathway of regulation of actin cytoskeleton should maintain the morphology of hair cells, and that noise will imbalance the pathway and lead to the degeneration of hair cells. If there are genes that can protect this pathway from noise-induced destruction, the hair cells can be protected, thereby protecting hearing.

In humans, hypoxia is an influential causative element of inner ear disease, and the role of HIF-1 in the regulation of oxygen homeostasis in the inner ear, such as regulation, energy supply, cell proliferation, or death is of interest. Insufficient blood supply after noise exposure leads to a decrease in the oxygen partial pressure, leaving the cochlea in a hypoxic state (pathology 2009). Hypoxic environmental preconditioning prevents noise-induced hearing loss in CBA/J and CBA/CAJ mice by upregulating HIF-1α in the organ of Corti (Gagnon et al. 2007). Another study proposed the use of cobalt chloride treatment, which up-regulates HIF-1α and protects hearing in noise-exposed mice (Chung et al. 2011). We suggest that the HIF-1 signaling pathway, which activates the transcription of diverse genes that enable cells to survive under hypoxic conditions, plays a crucial role in triggering protective metabolic changes in response to hypoxia (pathology 2009), resists noise, maintains cochlear homeostasis, and thus protects hearing.

The differential genes selected in the R_NE and the S_NE were due to different susceptibility of the mouse cochlea to noise; therefore, the enrichment pathways were associated with noise susceptibility. Among them, apoptosis and the NF-κB signaling pathway deserve our attention. Numerous studies have now demonstrated that cochlear hair cells can undergo apoptosis under noise exposure. Many drugs or methods have also been found to inhibit hair cell apoptosis to protect hearing. Therefore, we suggest that noise initiates the apoptosis pathway in hair cells, making the cochlea increasingly sensitive to noise.

While the NF-κB signaling pathway may be a defense pathway, it has been demonstrated that in the auditory system, the NF-κB signaling pathway can be activated by noise (Zhang et al. 2019a) and can prevent noise-induced hearing loss (Tahera et al. 2006). Investigations on the use of photobiomodulation (PBM) for noise-induced hearing loss found that PBM can activate NF-κB to protect the cochlea from oxidative stress and apoptosis (Tamura et al. 2016). In other studies, mice lacking the p50 subunit of NF-κB were found to have a higher sensitivity to noise exposure (Lang et al. 2006). Therefore, there is no doubt that the NF-κB signaling pathway can resist noise damage.

In this experiment, we found that ABR can be used to determine the presence of hearing loss. Otoacoustic emission can also be performed to detect cochlear amplification function and hair cell function integrity. In addition, there may have been asymmetric hearing loss, as we only tested the hearing threshold of one ear. Further experiments could be performed in the future to address these limitations.

## 5. Conclusions

This study revealed that mice with the same genetic background show different susceptibility to noise, and transcriptome specific changes were apparent after noise exposure. DEGs were found through variant analysis. Bioinformatic analysis revealed the functional implication of these genes. Changes in the transcriptome of these genes may affect cochlear susceptibility to noise. In the next step, the experimental results of this study should be verified using knockout mice. Noise-induced hearing loss is an irreversible disease; hence, its prevention is particularly important. Workers exposed to the same noise over the same occupational years experience varying degrees of noise-induced hearing loss. We can use gene therapy to prevent work-related injuries in these workers. As the society continues to evolve, people are becoming more susceptible to noise, which in turn damages their hearing. This study can help with future genetic screening, predict individual susceptibility to noise, and prevent noise-induced hearing loss using gene therapy.

## Data availability

The authors affirm that all data necessary for confirming the conclusions of the article are present within the article, figures, and tables. Supplementary material has been uploaded to https://gsajournals.figshare.com/.

## Acknowledgments

Haibo Wang, Zhaoming Fan, Lei Xu, and Jing Cai designed and supervised the project. Siyue Wang and Jing Cai performed experiments and acquired the data. Jing Cai and Ligang Kong analyzed and interpreted the results. Siyue Wang and Jing Cai wrote the manuscript. Xiuyue Biol (Jinan, China) analyzed the results of RNA sequencing.

## Funding

This study was supported by the National Natural Science Foundation of China (No. 82101226 and No. 81700918/H1304).

## Conflict of Interest

The authors declare no relevant conflict of interests.

## References

Beaulac HJ, Gilels F, Zhang J, Jeoung S, death PMWJC, disease. 2021. Primed to die: An investigation of the genetic mechanisms underlying noise-induced hearing loss and cochlear damage in homozygous foxo3-knockout mice. 12(7):682.

Chung JW, Shin J-E, Han KW, Ahn JH, Kim Y-J, Park J-W, toxicology H-SSJE, pharmacology. 2011. Up-regulation of hypoxia-inducible factor-1 alpha by cobalt chloride prevents hearing loss in noise-exposed mice. 31(1):153–159.

Davis RR, Newlander JK, Ling X, Cortopassi GA, Krieg EF, research LCEJH. 2001. Genetic basis for susceptibility to noise-induced hearing loss in mice. 155(1-2):82–90.

Ding T, Yan A, medicine KLJBjoh. 2019. What is noise-induced hearing loss? 80(9):525–529.

Du M-G, Liu F, Chang Y, Tong S, Liu W, Chen Y-J, chemistry PXJTJob. 2021. Neddylation modification of the u3 snorna-binding protein rrp9 by smurf1 promotes tumorigenesis. 297(5):101307.

Erway LC, Shiau YW, Davis RR, research EFKJH. 1996. Genetics of age-related hearing loss in mice. Iii. Susceptibility of inbred and f1 hybrid strains to noise-induced hearing loss. 93(1-2):181–187.

Fairfield DA, Lomax MI, Dootz GA, Chen S, Galecki AT, Benjamin IJ, Dolan DF, research RAAJJon. 2005. Heat shock factor 1-deficient mice exhibit decreased recovery of hearing following noise overstimulation. 81(4):589–596.

Feng Y, Pan T-C, Pant DK, Chakrabarti KR, Alvarez JV, Ruth JR, discovery LACJC. 2014. Spsb1 promotes breast cancer recurrence by potentiating c-met signaling. 4(7):790–803.

Fu X, Zhang L, Jin Y, Sun X, Zhang A, Wen Z, Zhou Y, Xia M, plasticity JGJN. 2016. Loss of myh14 increases susceptibility to noise-induced hearing loss in cba/caj mice. 2016:6720420.

Gagnon PM, Simmons DD, Bao J, Lei D, Ortmann AJ, research KKOJH. 2007. Temporal and genetic influences on protection against noise-induced hearing loss by hypoxic preconditioning in mice. 226(1-2):79–91.

Graham CE, Basappa J, disease DEVJNo. 2010. A corticotropin-releasing factor system expressed in the cochlea modulates hearing sensitivity and protects against noise-induced hearing loss. 38(2):246–258.

Gratton MA, Eleftheriadou A, Garcia J, Verduzco E, Martin GK, Lonsbury-Martin BL, research AEVJH. 2011. Noise-induced changes in gene expression in the cochleae of mice differing in their susceptibility to noise damage. 277(1-2):211–226.

Harding GW, Bohne BA, research JDVJH. 2005. The effect of an age-related hearing loss gene (ahl) on noise-induced hearing loss and cochlear damage from low-frequency noise. 204(1-2):90–100.

Henderson D, Subramaniam M, Ear FABJ, hearing. 1993. Individual susceptibility to noise-induced hearing loss: An old topic revisited. 14(3):152–168.

Holme RH, JARO KPSJJotAfRiO. 2004. Progressive hearing loss and increased susceptibility to noise-induced hearing loss in mice carrying a cdh23 but not a myo7a mutation. 5(1):66–79.

Honkura Y, Matsuo H, Murakami S, Sakiyama M, Mizutari K, Shiotani A, Yamamoto M, Morita I, Shinomiya N, Kawase T et al. 2016. Nrf2 is a key target for prevention of noise-induced hearing loss by reducing oxidative damage of cochlea. 6:19329.

Hu BH, Henderson D, research TMNJH. 2002. F-actin cleavage in apoptotic outer hair cells in chinchilla cochleas exposed to intense noise. 172(1-2):1–9.

Jamesdaniel S, Hu B, Kermany MH, Jiang H, Ding D, Coling D, proteomics RSJJo. 2011. Noise induced changes in the expression of p38/mapk signaling proteins in the sensory epithelium of the inner ear. 75(2):410–424.

Kim H-J, Kim HJ, Kim M-K, Bae MK, Sung HY, Ahn J-H, Kim YH, Kim SC, Biochemical WJJ, communications br. 2019. Spsb1 enhances ovarian cancer cell survival by destabilizing p21. 510(3):364–369.

Kozel PJ, Davis RR, Krieg EF, Shull GE, research LCEJH. 2002. Deficiency in plasma membrane calcium atpase isoform 2 increases susceptibility to noise-induced hearing loss in mice. 164(1-2):231–239.

Lang H, Schulte BA, Zhou D, Smythe N, Spicer SS, Neuroscience RASJTJontojotSf. 2006. Nuclear factor kappab deficiency is associated with auditory nerve degeneration and increased noise-induced hearing loss. 26(13):3541–3550.

Lavinsky J, Crow AL, Pan C, Wang J, Aaron KA, Ho MK, Li Q, Salehide P, Myint A, Monges-Hernadez M et al. 2015. Genome-wide association study identifies nox3 as a critical gene for susceptibility to noise-induced hearing loss. 11(4):e1005094.

Le TN, Straatman LV, Lea J, head BWJJoo-, cervico-faciale nsLJdo-r-ledc. 2017. Current insights in noise-induced hearing loss: A literature review of the underlying mechanism, pathophysiology, asymmetry, and management options. 46(1):41.

Liu J, Lv L, Gong J, Tan Y, Zhu Y, Dai Y, Pan X, Huen MSY, Li B, Tsao SW et al. 2018. Overexpression of f-box only protein 31 predicts poor prognosis and deregulates p38α- and jnk-mediated apoptosis in esophageal squamous cell carcinoma. 142(1):145–155.

Liu SY, Song WQ, Xin JR, Li Z, Lei S, Chen YQ, Zhao TY, Wang HY, Xu LW, Zhang MB et al. 2021. Nrn1 and cat gene polymorphisms, complex noise, and lifestyles interactively affect the risk of noise-induced hearing loss. 34(9):705–718.

Manne RK, Agrawal Y, Malonia SK, Banday S, Edachery S, Patel A, Kumar A, Shetty P, chemistry MKSJTJob. 2021. Fbxl20 promotes breast cancer malignancy by inhibiting apoptosis through degradation of puma and bax. 297(4):101253.

Mao H, plasticity YCJN. 2021. Noise-induced hearing loss: Updates on molecular targets and potential interventions. 2021:4784385.

Marioni JC, Mason CE, Mane SM, Stephens M, research YGJG. 2008. Rna-seq: An assessment of technical reproducibility and comparison with gene expression arrays. 18(9):1509–1517.

McGrath J, Roy P, cell BJPJSi, biology d. 2017. Stereocilia morphogenesis and maintenance through regulation of actin stability. 65:88–95.

Miao L, Ji J, Wan L, Zhang J, Yin L, science YPJE, international pr. 2019. An overview of research trends and genetic polymorphisms for noise-induced hearing loss from 2009 to 2018. 26(34):34754–34774.

Miao L, Zhang J, Yin L, research YPJIjoe, health p. 2021. Tmt-based quantitative proteomics reveals cochlear protein profile alterations in mice with noise-induced hearing loss. 19(1).

Ohlemiller KK, McFadden SL, Ding DL, Flood DG, Reaume AG, Hoffman EK, Scott RW, Wright JS, Putcha GV, Audiology RJSJ et al. 1999. Targeted deletion of the cytosolic cu/zn-superoxide dismutase gene (sod1) increases susceptibility to noise-induced hearing loss. 4(5):237–246.

Ohlemiller KK, McFadden SL, Ding DL, Lear PM, JARO YSHJJotAfRiO. 2000. Targeted mutation of the gene for cellular glutathione peroxidase (gpx1) increases noise-induced hearing loss in mice. 1(3):243–254.

oto-laryngologica HSLJA. 1992. Influence of genotype and age on acute acoustic trauma and recovery in cba/ca and c57bl/6j mice. 112(6):956–967.

Partearroyo T, Murillo-Cuesta S, Vallecillo N, Bermúdez-Muñoz JM, Rosa LR-dl, Mandruzzato G, Celaya AM, Zeisel SH, Pajares MA, Varela-Moreiras G et al. 2019. Betaine-homocysteine s-methyltransferase deficiency causes increased susceptibility to noise-induced hearing loss associated with plasma hyperhomocysteinemia. 33(5):5942–5956.

pathology XSJTAjo. 2009. Cochlear pericyte responses to acoustic trauma and the involvement of hypoxia-inducible factor-1alpha and vascular endothelial growth factor. 174(5):1692–1704.

Qin Y, discovery SSMJC. 2014. Spsb1 may have met its match during breast cancer recurrence. 4(7):760–761.

Raphael Y, neurology RAAJE. 1992. Early microfilament reorganization in injured auditory epithelia. 115(1):32–36.

Schick B, Praetorius M, Eigenthaler M, Jung V, Müller M, Walter U, Cell MKJ, research t. 2004. Increased noise sensitivity and altered inner ear mena distribution in vasp-/-mice. 318(3):493–502.

Shone G, Altschuler RA, Miller JM, research ALNJH. 1991. The effect of noise exposure on the aging ear. 56(1-2):173–178.

Sliwinska-Kowalska M, research MPJM. 2013. Contribution of genetic factors to noise-induced hearing loss: A human studies review. 752(1):61–65.

Tabuchi K, Suzuki M, Mizuno A, letters AHJN. 2005. Hearing impairment in trpv4 knockout mice. 382(3):304–308.

Tahera Y, Meltser I, Johansson P, Bian Z, Stierna P, Hansson AC, research BCJJon. 2006. Nf-kappab mediated glucocorticoid response in the inner ear after acoustic trauma. 83(6):1066–1076.

Tamura A, Matsunobu T, Tamura R, Kawauchi S, Sato S, research ASJB. 2016. Photobiomodulation rescues the cochlea from noise-induced hearing loss via upregulating nuclear factor κb expression in rats. 1646:467–474.

Themann CL, America EAMJTJotASo. 2019. Occupational noise exposure: A review of its effects, epidemiology, and impact with recommendations for reducing its burden. 146(5):3879.

Vlajkovic SM, Ambepitiya K, Barclay M, Boison D, Housley GD, research PRTJH. 2017. Adenosine receptors regulate susceptibility to noise-induced neural injury in the mouse cochlea and hearing loss. 345:43–51.

Waldhaus J, Cimerman J, Gohlke H, Ehrich M, Müller M, one HLJP. 2012. Stemness of the organ of corti relates to the epigenetic status of sox2 enhancers. 7(5):e36066.

Wang H, Xiao W, Zhou Q, Chen Y, Yang S, Sheng J, Yin Y, Fan J, research JZJC. 2009a. Bystin-like protein is upregulated in hepatocellular carcinoma and required for nucleologenesis in cancer cell proliferation. 19(10):1150–1164.

Wang Z, Gerstein M, Genetics MSJNr. 2009b. Rna-seq: A revolutionary tool for transcriptomics. 10(1):57–63.

Yan D, Zhu Y, Walsh T, Xie D, Yuan H, Sirmaci A, Fujikawa T, Wong ACY, Loh TL, D. L et al. 2013. Mutation of the atp-gated p2x(2) receptor leads to progressive hearing loss and increased susceptibility to noise. 110(6):2228–2233.

Yao Q, Wang H, Chen H, Li Z, Jiang Y, Li Z, Wang J, Xing Y, Liu F, Yu D et al. 2022. Essential role of sptan1 in cochlear hair cell morphology and function via focal adhesion signaling. 59(1):386–404.

Zhang G, Zheng H, Pyykko I, research JZJH. 2019a. The tlr-4/nf-κb signaling pathway activation in cochlear inflammation of rats with noise-induced hearing loss. 379:59–68.

Zhang L, Lin J, Rna KYJ. 2013. Structural and functional analysis of the u3 snorna binding protein rrp9. 19(5):701–711.

Zhang S, Ding E, Yin H, Zhang H, genomics BZJIjo. 2019b. Research and discussion on the relationships between noise-induced hearing loss and atp2b2 gene polymorphism. 2019:5048943.

Zhang X, Ni Y, Liu Y, Zhang L, Zhang M, Fang X, Yang Z, Wang Q, Li H, Xia Y et al. 2019c. Screening of noise-induced hearing loss (nihl)-associated snps and the assessment of its genetic susceptibility. 18(1):30.

Zhou J, Shi Z, Zhou L, Hu Y, open MZJB. 2020. Occupational noise-induced hearing loss in china: A systematic review and meta-analysis. 10(9):e039576.

Zhou X-X, Chen S, Xie L, Ji Y-Z, Wu X, Wang W-W, Yang Q, Yu J-T, Sun Y, Lin X et al. 2016. Reduced connexin26 in the mature cochlea increases susceptibility to noise-induced hearing lossin mice. 17(3):301.

